# Facilitating Replication and Reproducibility in Team Science: The ‘projects’ R Package

**DOI:** 10.1101/540542

**Authors:** Nikolas I. Krieger, Adam T. Perzynski, Jarrod E. Dalton

**Author notes:** Corresponding author: (NIK).

## Abstract

The contemporary scientific community places a growing emphasis on the reproducibility of research. The `projects` R package is a free, open-source endeavor created in the interest of facilitating reproducible research workflows. It adds to existing software tools for reproducible research and introduces several practical features that are helpful for scientists and their collaborative research teams. For each individual project, it supplies an intuitive framework for storing raw and cleaned study data sets, and provides script templates for protocol creation, data cleaning, data analysis and manuscript development. Internal databases of project and author information are generated and displayed, and manuscript title pages containing author lists and their affiliations are automatically generated from the internal database. File management tools allow teams to organize multiple projects. When used on a shared file system, multiple researchers can harmoniously contribute to the same project in a less punctuated manner, reducing the frequency of misunderstandings and the need for status updates.

## Introduction

The past few years have yielded much discussion and controversy in scientific circles surrounding the so-called replication crisis, in which results of important studies may fail to generalize to other samples. (1,2) Correspondingly, reproducibility and replication are increasingly central goals of the contemporary scientific process. Journals are increasingly emphasizing replicability into account when scrutinizing submissions, with some establishing replication standards for submitted articles. (3)

Reproducibility and replication are related but distinct. The reproducibility (or internal reproducibility) of a study is the ability of its readers to follow its methods and workflow that the researchers used to generate their reported result from their study data. Actively maintaining and archiving this workflow is important to the evaluation of the research: a study is internally reproducible when other researchers can follow the same workflow to achieve the same results from the same data. The external reproducibility of a study is the extent to which its methods and workflow can be meaningfully applied to external data (i.e., data that are different from the original study data). In turn, researchers who externally reproduce the same workflow to achieve similar results to those of the original study using different data provide evidence of replicability. (4) Thus, a study can be both internally and externally reproducible but not replicable if external reproduction attempts fail to produce results similar to those of the original study.

There exist today widely available tools that aid with reproducible research, such as R and other statistical programming languages, that allow for precise documentation of some of the most detail-oriented portions of a project workflow. Researchers can distribute their code scripts alongside their results in order to communicate the integrity of their data processing and analysis. Unfortunately, statistical programming languages per se only contribute to research reproducibility insofar as individual statistical programmers are able (i) to use these tools effectively and (ii) to integrate their own use of these tools with their collaborators’ work-which may not necessarily be oriented towards reproducibility.

The goal of the `projects` R package is to provide a set of tools that support an efficient project management workflow for statisticians and data scientists who perform reproducible research within team science environments. The `projects` package is built upon some existing tools for reproducible research, particularly RStudio, the R integrated development environment in which it dwells, and R Markdown, the file structure that allows users to assemble datasets, to perform analyses and to write manuscripts in a single file. The `projects` package is oriented towards efficient and reproducible academic research manuscript development and incorporates protocol and analysis templates based on widely-accepted reporting guidelines (viz., CONSORT and STROBE). When used on a shared file system (e.g., a server), the `projects` package provides infrastructure for collaborative research: multiple researchers can work on the same project and keep track of its progress without having to request updates.

The primary features of the `projects` R package are the following:

- Relational database containing details of projects, project coauthors and their affiliations, so that author details generally need to be entered only once;
- Tools for editing metadata associated with projects, authors and affiliations;
- Automated file structure supporting reproducible research workflow;
- Report templates that automatically generate title page headers, including a numbered author list and corresponding affiliations;
- Full RStudio integration via R Markdown, including customizable styling via cascading style sheets (CSS);
- Customization, including the ability to add and to edit templates for protocols and reports, and the ability to change default project directory and file structures; and
- Organization and management functionality, including the ability to group, archive and delete projects.

At its outset, the `projects` package creates a folder called */projects* in a user-specified location. This directory will contain all research projects created and maintained by the user. The */projects* folder will also contain a relational database of the research projects and the persons who contribute to them. The database allows for users to input important metadata about the projects and authors, including stage of research and contact information. Once this higher-level folder is created, users run R functions to create projects, each of which is given its own folder. New project folders automatically include aptly named subfolders and templates in order to guide the project workflow (e.g., a “data” subfolder; a “datawork” R Markdown template). Henceforth, users can begin working on the research project and edit as needed the metadata of the project itself and its authors. To lessen the burden of the mundane details of manuscript writing, the `projects` package can output lines to the console that, when copied into an R Markdown file, generates a title page with all relevant authorship information of any given project. Finally, since users may create dozens of projects over time, users can run functions to organize their projects within grouping subfolders of the main */projects* folder.

## Conceptual Framework: Reproducible Research Workflows

Although researchers of different disciplines may operate in nuanced ways, there are aspects of the project workflow that are common to most investigations. First, studies are conceptualized and designed according to a protocol that details the research questions and planned analyses. Data are collected, manipulated (or “tidied”) in order to make data analysis possible. The results of the analyses are compiled into a report, and ultimately an academic manuscript is drafted and submitted for wider distribution.

When navigating this workflow, researchers strive for reproducibility wherever possible but especially during the intermediate, data-focused phases of the workflow. While primary data sources for research projects can be complex, dynamic and diverse, a reproducible analytic workflow ultimately should incorporate a “frozen” dataset that reflects a given set of queries or data collection forms at a specific point in time. A frozen dataset represents the study data’s earliest state of simultaneous digitization and consolidation and is almost invariably a digital file or set of files that standard data analysis software can process (e.g., a comma-separated values, CSV, file; a series of CSV files; a compressed-format dataset in the researcher’s analytic programming language of choice).

From this point in the workflow through the reporting stage, total reproducibility is expected (see Fig 1). Thanks to modern data analysis via statistical programming languages, a reader should be able to exactly reproduce all data-derived results from the frozen dataset alone. Under the reproducible research framework, scientists should operate under the assumption that another scientist with access to the exact scripts the researchers used to produce their results should be able not only to regenerate the study results but to scrutinize every component of the analysis, beginning with data cleaning operations performed on the frozen dataset.

**Fig 1:**
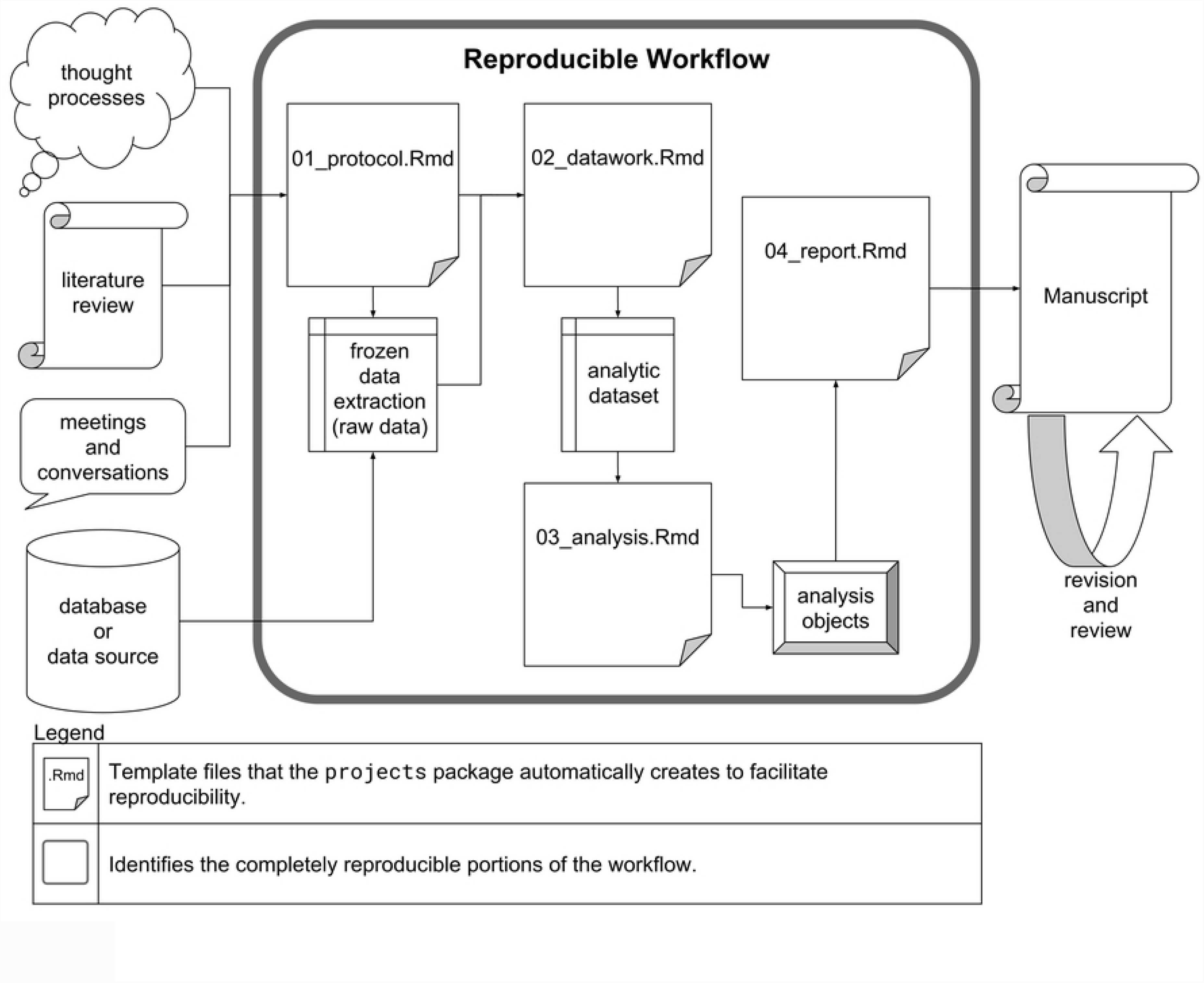
The entire workflow of manuscript creation with the reproducible portions encircled.

The middle stage of the assumed study workflow can be performed with near perfect reproducibility, but the beginning and ending stages may not. Researchers cannot document every thought process, literature probe and informal conversation that contributes to the development of the initial study protocol, but they should strive to document it as meticulously as possible. Databases tend to be dynamic such that a given analytic dataset is merely a snapshot in time. As for the final stages of project development, journals require that manuscripts adhere to specific and unique stylistic guidelines and that they be digitally submitted with file types that are not independently conducive to reproducibility (e.g., .*pdf*). For instance, even as RStudio supports the creation of submission-ready documents directly from frozen datasets, the vast majority of project teams include experts who do not use RStudio; therefore, the collaborative manuscript editing process ultimately takes place in an environment (e.g., Microsoft Word) that only supports total reproducibility with extraordinary effort. In light of these realities, during manuscript creation researchers must do their best, keeping the process in reproducible environments for as long as possible and otherwise documenting significant changes and alterations.

## projects R package

### Installation

The `projects` R package can be installed with:

~~~
**install.packages**("projects")
~~~

### Initial Setup

All projects that the user creates with the `projects` package-as well as its infrastructure-reside in a main folder called */projects*. Users need not manually create this directory, and in fact they are encouraged not to manually manipulate any folders that the `projects` package involves. Instead, users run the function `setup_ projects()`, providing the full file path of the directory in which the user wants the */projects* folder to reside.

### Interactive Metadata

Data about authors, institutional and/or department affiliations and projects are stored in .*rds* files within the main */projects* directory, so that the user only needs to enter these details once (unless, for example, a co-author changes their name or affiliations). These data are also used to assemble title pages of reports, with automatically generated author lists and lists of author affiliations. We provide a complete example of this process below in the **Demonstration** section.

The main metadata tables that the user interacts with are `projects()`, `authors()` and `affiliations()`, accessible via functions thusly named. The contents of these tables are described in Tables 1–3. Two additional tables are internally created to keep track of associations between authors and projects and between authors and affiliations (see **Internal Metadata Tables**).

**Table 1.** The projects() Metadata Table.

| Column | Description |
| --- | --- |
| `id` | An identification number, specifically an integer, unique among the other projects. This number can be used whenever needing to identify this project within `projects` package functions. |
| `title` | The title of the project. A nonambiguous substring of `title` (i.e., a substring that does not match any other project) can be used whenever needing to identify this project within `projects` package functions. This value is also printed in the YAML header of the “protocol” and “report” *.Rmd* templates that are automatically generated upon project creation. |
| `short_title` | An optional, unique nickname for the project. A nonambiguous substring of `short_title` (i.e., a substring that does not match any other project) can be used whenever needing to identify this project within `projects` package functions. This is useful if users cannot remember the long, formal project `title` nor the project `id`. |
| `current_owner` | The `id` of the author who is responsible for taking action in order that work on the project may proceed further. |
| `status` | A short description of the status of the project. For example, it may elaborate on the value of `current_owner` and/or `stage`. |
| `deadline_type` | A simple description of the meaning of the date contained in the next field, `deadline`. |
| `deadline` | A date indicating some kind of deadline whose meaning is described in the previous field, `deadline_type`. |
| stage | One of six predefined stages of project development that the project is currently in: `c("1: design", "2: data collection", "3: analysis", "4: manuscript", "5: under review", "6: accepted")`. |
| path | The full file path where the project folder is located. |
| corresp_auth | The id of the author who should be contacted for any correspondence relating to the project. This author's name will be specially marked on automatically generated title pages for this project, and his or her contact information will be specially displayed there as well in a "Corresponding Author" section. |
| creator | The id of the author who initially created the project, or the value of `Sys.info()["user"]` if the author who ran `new_project()` did not enter a value. |

**Table 2.**
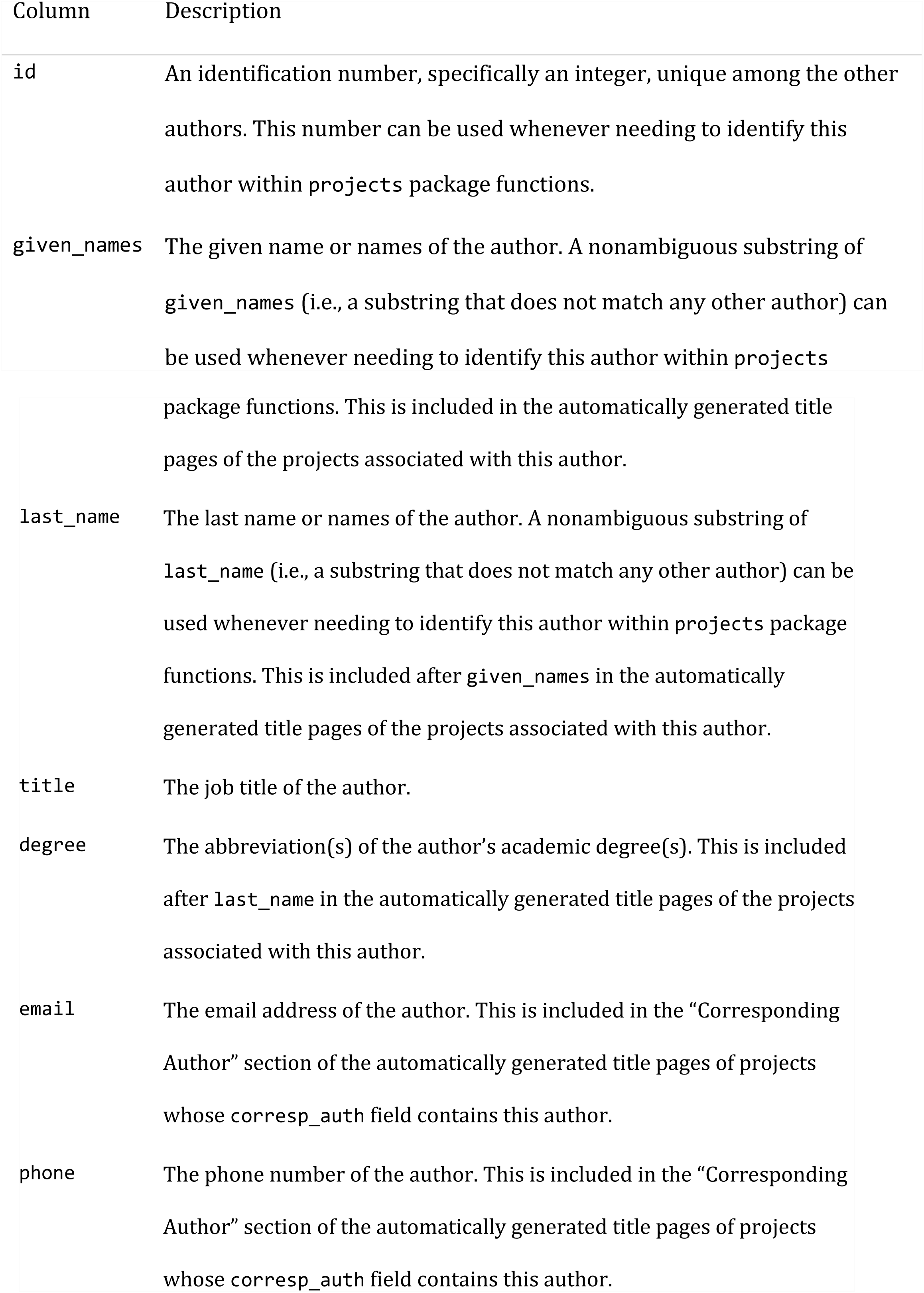
The authors() Metadata Table.

**Table 3.** The affiliations() Metadata Table.

| Column | Description |
| --- | --- |
| id | An identification number, specifically an integer, unique among the other affiliations. This number can be used whenever needing to identify this affiliation within projects package functions. |
| department_name | The department name of the affiliation. A nonambiguous substring of department_name (i.e., a substring that does not match any other affiliation) can be used whenever needing to identify this affiliation within projects package functions. This is included in the affiliations section of the automatically generated title page of projects associated with authors with this affiliation. |
| institution_name | The name of the overall institution of the affiliation. A nonambiguous substring of institution_name (i.e., a substring that does not match any other affiliation) can be used whenever needing to identify this affiliation within projects package functions. This is included after department_name in the affiliations section of the automatically generated title page of projects associated with authors with this affiliation. |
| address | The address of the affiliation. This is included after institution_name in the affiliations section of the automatically generated title page of projects associated with authors with this affiliation. It is also included in the “Corresponding Author” section of the title page when a project’s corresponding author has this affiliation as his or her primary (i.e., first) affiliation (see **Internal Metadata Tables**). |

### Internal Metadata Tables

In keeping with relational database theory, there are two .*rds* files that keep track of the many-to-many relationships between projects and authors and between authors and affiliations. Each has two columns, `id1` and `id2`, that contain the `id` numbers of these items. Each row of this table describes an association. Furthermore, the `projects` package keeps track of the order in which these associations appear so that the automatically generated title pages list authors and affiliations in the correct order. Users are able to run functions to reorder these associations as needed.

### Project File Structure

Users create individual project folders with the function `new_project()`. The name of each project folder is of the form p*XXXX*, where *XXXX* is the project’s `id` padded with 0s on the left side. When a project folder is created, it is automatically populated with folders and files as shown:

- */p*XXXX
  *–/data*
  *–/data_raw*
  *–/figures*
  *–/manuscript*
  *–/progs*
    - *01_protocol.Rmd*
    - *02_datawork.Rmd*
    - *03_analysis.Rmd*
    - *04_report.Rmd*
  *–pXXXX.Rproj*

The included subfolders serve to organize the project, while the .*Rmd* files are templates that facilitate the user’s workflow.

### File Management

The goal of the `projects` package is to provide a comprehensive set of tools managing project files in a way that is self-contained in R and independent of the underlying operating system. On a daily basis, researchers make, move, copy, delete and archive files. Through the `projects` package, researchers can perform all these actions in an organized manner with an automated file structure. In fact, users are advised not to manipulate the */projects* folder and its content with their operating system, so that the package does not lose track of these files. Multi-user application of `projects` requires a server or an otherwise shared directory where multiple users can access the */projects* folder. File-managing functions-along with all functions-are demonstrated below (see **Demonstration**).

### Other Features

The `projects` package supports style customization of manuscripts through cascading style sheets (CSS). When a project is created, a file called *style.css* is created alongside the *Rmd* files in the */progs* folder; users can customize their protocol and report by editing this file. Users can also create their own template files for the datawork, analysis and manuscript .*Rmd* files. Lastly, the user is given the option to make these .*Rmd* files BibTeX-ready for streamlined bibliography creation.

### Demonstration

Upon installation, the `projects` package must be set up using `setup_projects()`. The user is to input the file path of the directory wherein the */projects* folder is desired to be located.

~~~
**library**(projects)
**setup_projects**("C:/")
## "projects"folder created at
## C:/projects
##
## Add affiliations with new_affiliation(), then add authors with
## new_author(), then create projects with new_project()
~~~

As the message suggests, it is in the user’s best interest to add affiliations, followed by authors and projects.

~~~
**new_affiliation**(department_name = "Department of Physics",
               institution_name = "University of North Science",
               address = "314 Newton Blvd, Springfield CT 06003")

## # A tibble: 1 x 4
##     id department_name  institution_name  address
##   <int> <chr>             <chr>            <chr>
## 1   1 Department of Phys~ University of North S~ 314 Newton Blvd, Sprin~
~~~

This affiliation has been successfully added to the “affiliations” table in the `projects` relational database. The next code chunk creates a few more affiliations (output not included).

~~~
**new_affiliation**(department_name = "Impossibles Investigation Team",
              institution_name = "Creekshirebrook Academy of Thinks",
              address = "Let Gade 27182, 1566 Copenhagen, Denmark")
**new_affiliation**(department_name = "Statistical Consulting Unit",
        institution_name = "Creekshirebrook Academy of Thinks",
        address = "196 Normal Ave, Columbus, OH ", id = 50)
~~~

Note that we chose a specific `id` number (50) for the affiliation called the “Impossibles Investigation Team.”

Now we are ready to add authors to the “authors” table of the `projects` database.

~~~
**new_author**(given_names = "Scott", last_name = "Bug", title = "Professor",
            affiliations = **c**(2, "Physics"), degree = "PhD",
            email = "scottbug@imPOSSible.net", phone = "965-555-5556")
## New author:
## # A tibble: 1 x 7
##       id given_names   last_name   title   degree email    phone
##    <int><  chr>            <chr>    <chr>  <chr>      <chr>  <chr>
## 1      1 Scott           Bug     Profess~ PhD     scottbug@impossib~ 965-555-~
##

## # A tibble: 2 x 4
##   affiliation_id department_name             institution_name     address
##          <int>    <chr>                         <chr>         <chr>
## 1          2 Impossibles Invest~ Creekshirebrook Acad~ Let Gade 27182~
## 2          1 Department of Phys~ University of North ~ 314 Newton Blv~
~~~

In creating associations between Scott Bug and his affiliations, we were able to enter both the `id` number of one of them (2) and a substring of the `department_name` of the other (“Physics”). The email address was coerced to lowercase.

Now we will add more authors (output not included).

~~~
**new_author**(given_names = "Marie", last_name = "Curie", title = "Chemist",
       affiliations = "Unit", phone = "553-867-5309", id = 86)
**new_author**(given_names = "George Washington", last_name = "Carver",
       title = "Botanist", degree = "MS",
       affiliations = **c**(1, 2, 50), id = 1337)
**new_author**(given_name = "Archimedes", title = "Mathematician")
**new_author(last_name = "Wu", given_names = "Chien-Shiung",**
       title = "Physicist",
       affiliations = **c**("of North", "Statistical Consulting"),
       degree = "PhD", email = "wu@WU.wU")
~~~

Now that some authors and affiliations have been created, we can view these tables:

~~~
**authors**()

## # A tibble: 5 x 7
##     id given_names        last_name title       degree email       phone
##       <int> <chr>        <chr> <chr>           <chr> <chr>          <chr>
## 1    1 Scott    Bug    Professor   PhD   scottbug@impo~  965-555~
## 2    2 Archimedes  <NA>    Mathemat~   <NA> <NA>    <NA>
## 3 3 Chien-Shiung Wu Physicist PhD wu@wu.wu <NA>
## 4 86 Marie Curie Chemist <NA> <NA> 553-867~
## 5 1337 George Washing~ Carver Botanist MS <NA> <NA>

**affiliations**()

## # A tibble: 3 x 4
## id department_name institution_name address
## <int> <chr> <chr> <chr>
## 1 1 Department of Physics University of North Sc~ 314 Newton Blvd, Sp~
## 2 2 Impossibles Investig~ Creekshirebrook Academ~ Let Gade 27182, 156~
## 3 50 Statistical Consulti~ Creekshirebrook Academ~ "196 Normal Ave, Co~
~~~

Now we will showcase project creation:

~~~
**new_project**(title = "Achieving Cold Fusion", short_title = "ACF",
            authors = **c**("Bug", "Chien-Shiung", 86, 1337), current_owner = "Carver", corresp_auth = "Bug",
            stage = "1: design",
            deadline_type = "Pilot study", deadline = "2020-12-31",
            use_bib = TRUE)
##
## Project 1 has been created at
## C:/projects/p0001

## # A tibble: 1 x 8
## id title   short_title status   deadline_type deadline stage path
## <int> <chr>   <chr> <chr> <chr> <date> <fct>   <chr>
## 1   1 Achievi~ ACF   just c~ Pilot study   2020-12-31 1: d~ C:/pr~
##
## New project’s authors:

## # A tibble: 4 x 7
## author_id given_names last_name title degree email phone
## <int>      <chr>     <chr>    <chr>  <chr>  <chr>  <chr>
## 1     1     Scott     Bug     Profes~ PhD scottbug@imp~ 965-555~
## 2     3     Chien-Shiung     Wu     Physic~ PhD wu@wu.wu <NA>
## 3     86    Marie     Curie     Chemist <NA> <NA> 553-867~
## 4     1337  George Washin~     Carver     Botani~ MS <NA> <NA>
##
## Current owner:

## # A tibble: 1 x 7
## id given_names     last_name title     degree email phone
## <int> <chr>      <chr>      <chr> <chr>      <chr>      <chr>
## 1 1337 George Washington Carver Botanist MS <NA> <NA>

##
## Corresponding author:

## # A tibble: 1 x 7
##   id given_names last_name title degree email phone
## <int> <chr>   <chr>   <chr>   <chr>   <chr>  <chr>
## 1      1 Scott      Bug      Profess∼      PhD      scottbug@impossib∼      965-555-∼
##
## Creator: nkrieger
~~~

Since a creator was not specified, this field was populated with the value of

~~~
Sys.info()["user"].
~~~

Among other files and folders, this line of code created the files *01*_*protocol.Rmd* and *04*_*report.Rmd*, which both include code to create a title page exhibiting a preformatted list of authors and their affiliations.

Here is what the top of these files look like:

~~~
---
title: "Achieving Cold Fusion"
output:
 word_document: default
 html_document:
 css: style.css
bibliography: p0001.bib
---
**_Scott Bug, PhD;^1,2^\* Chien-Shiung Wu, PhD;^2,3^ Marie Curie;^3^ and George Washington Carver, MS^1,2,3^_**
| ^1^ Impossibles Investigation Team, Creekshirebrook Academy of Thinks, Let Gade 27182, 1566 Copenhagen, Denmark
| ^2^ Department of Physics, University of North Science, 314 Newton Blvd, Springfield CT 06003
| ^3^ Statistical Consulting Unit, Creekshirebrook Academy of Thinks, 196 Normal Ave, Columbus, OH
|
| \* Corresponding author
| Let Gade 27182, 1566 Copenhagen, Denmark
| 965-555-5556
| scottbug@impossible.net
|
| Funding:
\pagebreak
~~~

We notice that the author order given to the `authors` argument in `new_project()` command has been preserved; furthermore, Scott Bug has been marked as the corresponding author and his contact information has been included. Once this. *Rmd* file is rendered, this will become a proper title page.

More projects can be created as follows:

~~~
**new_project**(title = "Weighing the Crown", short_title = "Eureka!",
         authors = "Archimedes", current_owner = "Archimedes",
         corresp_auth = "Archimedes", stage = 6)
**new_project**(title = "How I Learned to Stop Worrying and Love the Bomb",
        short_title = "Dr. Strangelove", authors = **c**("wu", 1),
        creator = "wu", current_owner = "George",
        corresp_auth = "George",stage = "under review",
        deadline_type = "2nd revision",deadline = "2030-10-8", id = 1945,
        status = "debating leadership changes", path = "top_secret",
        make_directories = TRUE)
**new_project**(title = "Understanding Radon", short_title = "Rn86",
        authors = 86, creator = 86, corresp_auth = 86, stage = "3",
        status = "Safety procedures", protocol = "CONSORT",
        use_bib = TRUE)
~~~

### Below is the list of all projects that have been created

~~~
**projects**()
## # A tibble: 4 x 11
## id title short_title current_owner status deadline_type deadline
## <int> <chr> <chr> <int> <chr> <chr> <date>
## 1 1 Achi∼ ACF 1337 just ∼ Pilot study 2020-12-31
## 2 2 Weig∼ Eureka! 2 just ∼ <NA> NA
## 3 3 Unde∼ Rn86 86 Safet∼ <NA> NA
## 4 1945 How ∼ Dr. Strang∼ 1337 debat∼ 2nd revision 2030-10-08
## # … with 4 more variables: stage <fct>, path <chr>, corresp_auth <int>,
## # creator <chr>
~~~

Projects, authors and affiliations can be edited with edit_project(), edit_ author() and edit_affiliation(), respectively. For example, we can add affiliations to and remove affiliations from an author with:

~~~
**edit_author**(author = "Bug", affiliations = **∼ +** 50 **-** impossibles)
## Edited author:
## # A tibble: 1 x 7
## id given_names last_name title degree email phone
## <int> <chr> <chr> <chr> <chr> <chr> <chr>
## 1 1 Scott Bug Profess∼ PhD scottbug@impossib∼ 965-555-∼
##
## Edited author’s affiliations:
## # A tibble: 2 x 4
## affiliation_id department_name institution_name address
## <int> <chr> <chr> <chr>
## 1 1 Department of Phys ∼ University of North ∼ 314 Newton Blv ∼
## 2 50 Statistical Consul ∼ Creekshirebrook Acad ∼ "196 Normal Av ∼
~~~

When adding or removing affiliations/authors from an author/project, a one-sided `formula`is used: it must begin with a tilde (`∼`), and elements are added with + and removed with `-`. Elements can be referred to by their `id` numbers or their names, as described above.

A formula is also used in the `authors` argument in `edit_project()`

~~~
**edit_project**("Cold", title = "Cold Fusion Is Actually Impossible",authors = **∼** "archi", stage = "accepted")
## Edited project info:

## # A tibble: 1 x 11
## 1 id title short_title current_owner status deadline_type deadline
## <int> <chr> <chr> <int> <chr> <chr> <date>
## 1 Cold∼ ACF 1337 just ∼ Pilot study 2020-12-31
## # … with 4 more variables: stage <fct>,path <chr>, corresp_auth <int>,
## # creator <chr>
##
## Edited project’s authors:
## # A tibble: 5 x 7
## author_id given_names last_name title degree email phone
## <int> <chr> <chr> <chr> <chr> <chr> <chr>
## 1 1 Scott Bug Professor PhD scottbug@im∼ 965-55∼
## 2 3 Chien-Shiung Wu Physicist PhD wu@wu.wu <NA>
## 3 86 Marie Curie Chemist <NA> <NA> 553-86∼
## 4 2 Archimedes <NA> Mathemat∼ <NA> <NA> <NA>
## 5 1337 George Washin∼ Carver Botanist MS <NA> <NA>
##
## title: "Cold Fusion Is Actually Impossible"
##
##
## **_Scott Bug, PhD;^1,2^\* Chien-Shiung Wu, PhD;^1,2^ Marie Curie;^2^ Archimedes; and George Washington Carver, MS^1,2,3^_**
##
## | ^1^ Department of Physics, University of North Science, 314 Newton Blvd, Springfield CT 06003
## | ^2^ Statistical Consulting Unit, Creekshirebrook Academy of Thinks, 196 Normal Ave, Columbus, OH
## | ^3^ Impossibles Investigation Team, Creekshirebrook Academy of Thinks,
Let Gade 27182, 1566 Copenhagen, Denmark
## |
## | \* Corresponding author
## | 314 Newton Blvd, Springfield CT 06003
## | 965-555-5556
## | scottbug@impossible.net ## |
## | Funding:
~~~

Here, the `title` and `stage` of the project have also been edited. The default behavior of `edit_project()` is to reprint the project `title` as well as the other elements of the title page. The user can then copy and paste this header into the *01_protocol.Rmd* and *04_report.Rmd* files.

We also notice that the default behavior when adding elements is to place them before the last author (unless there was only one author). This occurs **after** elements are removed, as specified by any minus signs (`-`) in the `formula`.

Another function that affects author order and whose default behavior reprints project title page information is `reorder_authors()`:

~~~
**reorder_authors**(project = "Cold Fusion", "George", "Bug", 86)
## project info:
## # A tibble: 1 x 11
## id title short_title current_owner status deadline_type deadline
## <int> <chr> <chr> <int> <chr> <chr> <date>
## 1 1 Cold∼ ACF 1337 just ∼ Pilot study 2020-12-31
## # … with 4 more variables: stage <fct>, path <chr>, corresp_auth <int>,
## # creator <chr>
##
## Reordered project authors:
## # A tibble: 5 x 7
## author_id given_names last_name title degree email phone
## <int> <chr> <chr> <chr> <chr> <chr> <chr>
## 1 1337 George Washin∼ Carver Botanist MS <NA> <NA>
## 2 1 Scott Bug Professor PhD scottbug@im∼ 965-55∼
## 3 86 Marie Curie Chemist <NA> <NA> 553-86∼
## 4 3 Chien-Shiung Wu Physicist PhD wu@wu.wu <NA>
## 5 2 Archimedes <NA> Mathemat∼ <NA> <NA> <NA>
##
## title: "Cold Fusion Is Actually Impossible"
##
##
## **_George Washington Carver, MS;^1,2,3^ Scott Bug, PhD;^1,3^\* Marie Curie;^3^ Chien-Shiung Wu, PhD;^1,3^ and Archimedes_**
##
## | ^1^ Department of Physics, University of North Science, 314 Newton Blvd, Springfield CT 06003
## | ^2^ Impossibles Investigation Team, Creekshirebrook Academy of Thinks, Let Gade 27182, 1566 Copenhagen, Denmark
## | ^3^ Statistical Consulting Unit, Creekshirebrook Academy of Thinks, 196 Normal Ave, Columbus, OH
## |
## | \* Corresponding author
## | 314 Newton Blvd, Springfield CT 06003
## | 965-555-5556
## | scottbug@impossible.net
## |
## | Funding:
~~~

Importantly, `edit_author()`, `edit_affiliation()`, and `reorder_affiliations()` do **not** reprint the title pages of all affected projects. Fortunately, the user can reprint updated title pages with the `header()` function:

~~~
**edit_affiliation**(affiliation = "Impossibles",
     department_name = "Pseudoscience Debunking Unit")
**header**(project = "Cold")
title: "Cold Fusion Is Actually Impossible"
**_George Washington Carver, MS;^1,2,3^ Scott Bug, PhD;^1,3^\* Marie Curie;^3^ Chien-Shiung Wu, PhD;^1,3^ and Archimedes_**
| ^1^ Department of Physics, University of North Science, 314 Newton Blvd, Springfield CT 06003
| ^2^ Pseudoscience Debunking Unit, Creekshirebrook Academy of Thinks, Let Gade 27182, 1566 Copenhagen, Denmark
| ^3^ Statistical Consulting Unit, Creekshirebrook Academy of Thinks, 196 Normal Ave, Columbus, OH
|
| \* Corresponding author
| 314 Newton Blvd, Springfield CT 06003
| 965-555-5556
| scottbug@impossible.net
|
| Funding:
~~~

In order to organize projects, users can create subdirectories within the main */projects* folder where individual project folders can dwell. Among the examples above, this has already occurred with the project with the nickname (i.e., `short_title`) “Dr. Strangelove” because on its creation the arguments `path = top_secret` and `make_directories = TRUE` were included. The latter argument must be `TRUE` if the desired path does not already exist. Observe the `path` column among the existing projects:

**projects**() **%>% select**(id, short_title, path)

~~~
## # A tibble: 4 x 3
## id short_title path
## <int> <chr> <chr>
## 1 1 ACF C:/projects/p0001
## 2 Eureka! C:/projects/p0002
## 3 Rn86 C:/projects/p0003
## 4 1945 Dr. Strangelove C:/projects/top_secret/p1945
~~~

Users can also create subdirectories with the function `new_project_group()`:

~~~
**new_project_group**("Greek_studies/ancient_studies")
##
## The following directory was created:
## C:/projects/Greek_studies/ancient_studies
~~~

If a project has already been created, it can be moved **not** with `edit_project()` but with `move_project()`. Users can also copy projects using `copy_project()`; everything in the copy will be the same except its `id`, folder name (which, again, is based on its `id`), `path` (which, again, is based on its folder name), and the name of its.*Rproj* file (which has the same name as the folder).

~~~
**move_project**("Crown", path = "Greek_studies/ancient_studies")
**copy_project**(project_to_copy = "Radon",
path = "dangerous_studies/radioactive_studies/radon_studies", make_directories = TRUE)
**projects**(**c**("Crown", "Radon")) **%>% select**(id, title, path)
## # A tibble: 3 x 3
## id title path
## <int> <chr> <chr>
## 1 2 Weighing the C∼ C:/projects/Greek_studies/ancient_studies/p0002
## 2 3 Understanding ∼ C:/projects/p0003
## 3 4 Understanding ∼ C:/projects/dangerous_studies/radioactive_studies∼
~~~

Projects can also be archived; they are moved into a subdirectory called */archive* that is at the same level as the project folder (*/p*XXXX) before it was run. If this */archive* folder does not exist, it will be created.

~~~
**archive_project**("Strangelove")
## # A tibble: 1 x 11
## id title short_title current_owner status deadline_type deadline
## <int> <chr> <chr> <int> <chr> <chr> <date>
## 1 1945 How ∼ Dr. Strang∼ 1337 debat∼ 2nd revision 2030-10-08
<int> <chr>
## # … with 4 more variables: stage <fct>, path <chr>, corresp_auth <int>,
## # <chr>
## The above project was archived and has the file path
## C:/projects/top_secret/archive/p1945
~~~

When a project is archived, it is no longer included in `projects()` output unless the user sets `archived = TRUE`.

~~~
**projects**() **%>% select**(id, short_title, path)
## # A tibble: 4 x 3
## id short_title path
## <int> <chr> <chr>
## 1 1 ACF C:/projects/p0001
## 2 2 Eureka! C:/projects/Greek_studies/ancient_studies/p0002
## 3 3 Rn86 C:/projects/p0003
## 4 4 Rn86 C:/projects/dangerous_studies/radioactive_studies/rad∼

**projects**(archived = TRUE) **%>% select**(id, short_title, path)

## # A tibble: 5 x 3
## id short_title path
## <int> <chr> <chr>
## 1 1 ACF C:/projects/p0001
## 2 2 Eureka! C:/projects/Greek_studies/ancient_studies/p0002
## 3 3 Rn86 C:/projects/p0003
## 4 4 Rn86 C:/projects/dangerous_studies/radioactive_studies/rad∼
## 5 1945 Dr. strongelo∼ C:/projects/top_secret/archive/p1945
~~~

Lastly, affiliations, authors and projects can be deleted with `delete_affiliation()`, `delete_author()` and `delete_project()`, respectively. Deleting an author is complete: doing so removes the author from the `creator`, `current_owner` and `corresp_auth` fields of all projects. Furthermore, deleting a project also deletes the entire project folder. Users should use the `delete` functions with caution.

~~~
**delete_affiliation**(“north science”)
## # A tibble: 1 × 4
## id department_name institution_name address
## 1 <int> <chr> <chr> <chr>
## 1 1 Department of Phys∼ University of North S∼ 314 Newton Blvd, Sprin∼
## # A tibble: 1 x 4
## id department_name institution_name address
## <int> <chr> <chr> <chr>
## 1 <int> <chr> <chr> <chr>
## 1 1 Department of Phys∼ University of North S∼ 314 Newton Blvd, Sprin∼
## The above affiliation was deleted.
**delete_author**(2)
## # A tibble: 1 x 7
## id given_names last_name title degree email phone
## <int> <chr> <chr> <chr> <chr> <chr> <chr>
## 1 2 Archimedes <NA> Mathematician <NA> <NA> <NA>
## # A tibble: 1 x 7
## id given_names last_name title degree email phone
## <int> <chr> <chr> <chr> <chr> <chr> <chr>
## 1 2 Archimedes <NA> Mathematician <NA> <NA> <NA>
## The above author was deleted.
**delete_project**(“Crown”)
## # A tibble: 1 x 11
## id title short_title current_owner status deadline_type deadline
## <int> <chr> <chr> <int> <chr> <chr> <date>
## 1 2 Weig∼ Eureka! NA just ∼ <NA> NA
## # … with 4 more variables: stage <fct>, path <chr>, corresp_auth <int>, ## # creator <chr>
## # A tibble: 1 x 11
## id title short_title current_owner status deadline_type deadline
## <int> <chr> <chr> <int> <chr> <chr> <date>
## 1 2 Weig∼ Eureka! NA just ∼ <NA> NA
## # … with 4 more variables: stage <fct>, path <chr>, corresp_auth <int>,
## # creator <chr>
##
## The above project was deleted.
~~~

## Discussion

The `projects` package provides a comprehensive set of tools for reproducible team science workflows. Efficiency in project management, including manuscript development, is facilitated by an internal database that keeps record of project details as well as team members’ affiliations and contact information. For manuscripts, title pages are automatically generated from this database, and a selection of manuscript outlines compliant with reporting guidelines are available upon installation. Other workflow-related R packages exist, such as `workflowr`, which offers collaborative project file management on Github. The `projects` package, however, is unique in its focus on manuscript development within the typical academic research setting where data are firewalled and research teams collaborate within secure networks. (5)

The `projects` package builds upon existing tools that facilitate reproducibility in small ways. R Markdown, for example, allows reviewers to intuitively investigate R scripts piece by piece in order to inspect the integrity of data management and to validate intermediate results; however, the R Markdown file format does not by itself organize project files. In addition, the `here` package makes file paths generalizable so that networks of R scripts will run without error on multiple computers with different parent file structures; however, it does not standardize file names and paths in the project’s immediate working directory. The `projects` package not only adopts the benefits of R Markdown and `here` in its templates but also facilitates file management with a standardized file structure. Thus, the user’s network of R scripts and data objects maintains continuity. Collaborators and reviewers can—at least in principle—run a `projects` user’s entire workflow on their own computers without file-path-related errors. This streamlines both replication attempts as well as the investigation of reproducibility.

The main limitation of the `projects` package lies in its setting: it is confined to the R statistical programming language, which not all researchers know and use. Prospective users of the package who do not know R must spend time learning how to use it, and this drawback is compounded as the size of the transitioning research team increases. Researchers who do know R may need to spend time gaining proficiency with R Markdown and other `projects` package dependencies. In future work, we will explore ways to integrate the functionality of the projects package with other statistical programming languages (e.g., Python and SAS).

In spite of the inevitable learning curve that is present when adapting to a new programming language, the `projects` package is intuitive to use among regular users of R and R Markdown. We believe that the `projects` package may be useful for teams that manage multiple collaborative research projects in various stages of development. It has the potential to improve both the quality and efficiency of individual and team researchers while also rendering the task of maintaining reproducibility less cumbersome. The open-source nature of the R environment ensures that the `projects` package will only improve with time and use, as the scientific community continues to embrace the tools essential for maintaining a reproducible workflow.

## Acknowledgements

The authors of this package acknowledge the support provided by members of the Northeast Ohio Cohort for Atherosclerotic Risk Estimation (NEOCARE) investigative team: Claudia Coulton, Douglas Gunzler, Darcy Freedman, Neal Dawson, Michael Rothberg, David Zidar, David Kaelber, Douglas Einstadter, Alex Milinovich, Monica Webb Hooper, Kristen Hassmiller-Lich, Ye Tian (Devin), Kristen Berg, and Sandy Andrukat.

